# Fabrication of microcompartments with controlled size and shape for encapsulating active matter

**DOI:** 10.1101/2025.04.23.650156

**Authors:** Benoit Vianay, Christophe Guérin, Laurène Gressin, Magali Orhant-Prioux, Laurent Blanchoin, Manuel Théry, Alexandra Colin

## Abstract

In all living systems, the cytoplasm is separated from the external environment by membranes. This confinement imposes spatial constraints on the self-organization of internal components, filaments and organelles. While reconstituted systems are instrumental for understanding fundamental biological principles, traditional experiments often utilize volumes vastly larger than actual cells. In recent studies, water-in-oil droplets or giant unilamellar vesicles have been widely used to impose confinement. However, these compartments present imaging challenges and make precise protein content control difficult. To address these limitations, we have developed versatile microwells that are straightforward to implement, compatible with different types of imaging, suitable for long-term experiments, and capable of generating large amounts of data. These microwells are compatible with several surface treatments and a wide range of experimental techniques (such as micropatterning) making them a powerful tool for answering key questions in cell biology. We present here a detailed protocol of the fabrication of the microwells as well as characterization of the method to ensure quality throughout the manufacturing process. These microwells support various cytoskeleton-based processes including actin polymerization, dynamic steady-state actin networks, and composite actin-microtubule networks. More broadly, they can be used to encapsulate and study over time any kind of active matter.

## Introduction

In living systems, cell membranes create boundaries that isolate internal contents from the external environment. This compartmentalization establishes boundary conditions crucial for biological processes like morphogenesis for example. Beyond simple isolation, these enclosed environments and their boundaries serve dual purposes - they generate heterogeneity and permit specific biological processes to function in isolation from one another (Alberts *et al*, 2002).

Reconstituted systems are of primary importance in defining the minimum principles of complex biological behavior. However, most reconstitution experiments are conducted using volumes significantly larger than those found in actual cells. This prevents the observation or control of depletion effects that naturally occur within the constrained volumes of cells. Recognizing this limitation, research teams over the past decade have focused on developing methods to impose confinement on reconstituted systems, particularly those involving cytoskeletal components (Jia & Schwille, 2019; Bashirzadeh & Liu, 2019; Groaz *et al*, 2021; Fanalista *et al*, 2019; Kandiyoth & Michelot, 2023).

This research direction has been particularly followed in the field of synthetic cells, in which the compartment is of paramount importance for reconstituting the building blocks towards the first synthetic cell (Adamala *et al*, 2024; Schwille *et al*, 2018). In particular, water-in-oil droplets or giant unilamellar vesicles (GUVs) have emerged as preferred compartmentalization methods because these structures can deform (making them suitable for reconstituting cell division), or move (making them suitable for reconstituting motility) (Siton-Mendelson & Bernheim-Groswasser, 2016; Litschel & Schwille, 2021). However, these objects present significant imaging challenges as three-dimensional structures, typically requiring confocal microscopy for proper visualization. In addition, the amount of protein inside droplets or GUVs can be difficult to control precisely. This is why we decided to develop microwells that can be easily observed with several types of microscopies and in which the number and concentration of proteins can be well controlled over long periods of time (hours).

Microchambers have long been used to spatially constrain cells or reconstituted systems using a variety of materials and fabrication techniques (Manzoor *et al*, 2021). For example, they have been widely used to manipulate the shape of starfish embryos, yeast or human mesenchymal cells, making it possible to observe how changes in shape affect the position of the division plane, cytoskeleton assembly, or differentiation (Minc *et al*, 2011, 2009; Bao *et al*, 2017, 2018). In tissues, cells are organized in three dimensions; in particular, the development of microchambers has made it possible to mimic the cell niche and study the effect of cell-cell interaction on cell polarization (Bessy *et al*, 2021; Candelas *et al*, 2024).

For reconstituted systems, microchambers have mainly been used to impose mechanical constraints. For example, they have been used to place functional proteins on the sidewalls of microwells to spatially control microtubule gliding (Romet-Lemonne *et al*, 2005) and to study the positioning of microtubule asters (Laan *et al*, 2012). On the other hand, (Soares e Silva *et al*, 2011; Deshpande & Pfohl, 2015) have used microchambers to study actin bundles formation in various types of confinement. Microchambers have also been used to investigate how confinement dimensions influence the formation of active nematic phases in microtubule systems (Opathalage et al, 2019).

Based on these examples, confinement in reconstituted cytoskeleton-based systems has primarily been utilized to impose mechanical constraints on actin and microtubule networks. However, confinement also serves to limit the quantity of components, more closely mimicking cellular conditions.

In cell-sized confinement, Xenopus egg extracts encapsulated in water-in-oil droplets revealed scaling mechanisms of mitotic spindles by limiting available components (Good et al, 2013; Hazel et al, 2013). Despite this breakthrough, few studies have explored how component limitations affect the formation and maintenance of dynamic structures or evaluated the biochemical feedback mechanisms at play in biological processes. However, recent work demonstrated the importance of recycling processes for maintaining long-lived dynamic structures under limited component conditions (Colin et al, 2023a). In a subsequent study, protein turnover was demonstrated to be of primary importance for the coexistence of competitive actin networks when components are limited in quantity (Guérin et al, 2025).

Here, we present a detailed protocol for the manufacturing process of NOA microwells, which are well suited for cytoskeleton reconstitution on long timescales (several hours). We chose NOA because it is an inert material with glass-like properties, easy to handle and polymerize. Process characterization is provided throughout the microfabrication steps to ensure high microwell quality. These microwells were used to reconstitute cytoskeleton-based process in presence of a limited amount of proteins and mechanical constraints (Yamamoto *et al*, 2022; Colin *et al*, 2023a; Guérin *et al*, 2025; Sciortino *et al*, 2025) but they could be widely used to impose closed boundary conditions for various studies of cytoskeleton-based processes, particularly to vary compartment size and biochemical content and study their impact on the dynamics and overall organization of diverse structures.

## Materials and equipment

### 1. Microfabrication of SU8 and PDMS molds in order to build the microwells

#### 1.1. SU8 mold fabrication

##### 1.1.a. Material

Glass wafers

Adhesion promoter (Ti Prime, MicroChemicals)

SU8 3000 serie

SU8 Developper

Isopropanol 99.9% (34863, Sigma, CAS 67-63-0)

##### 1.1.b. Equipment

Mask (chrome or plastic) with the design and UV Lamp (UV KUB2, KLOE) OR Primo (Alveole) device

Oven or heating plate (from 60°C to 150°C)

Spin coater (WS-400BX-6NPP, Laurell)

#### 1.2. Primary PDMS mold, epoxy mold and PDMS pillars fabrication

##### 1.2.a. Material

Trichloro (1H,1H,2H,2H, perfluorooctyl) silane (448931, Sigma, CAS 78560-45-9)

PDMS (SYLGARD 184 silicone elastomer kit, Dow)

Glass petri dishes (diameter ~ 50 mm)

Epoxy resin (type R123, Bisphenol A/F epoxy resin) and hardener (type R614) from Soloplast-Vosschemie (Saint-Egreve, France)

##### 1.2.b. Equipment

Vacuum bell

Vacuum pump

Oven or heating plate (70°C to 150°C)

Centrifuge (for 50 mL Falcon tube, 500g)

#### 1.3. NOA microwells preparation

##### 1.3.a. Material

Coverglasses 20*20 mm n°1.5 (Agar Scientific L46520-15)

Ethanol 96% (UN1170 Carlo Erba, CAS 64-17-5)

Hellmanex (2% in water) (Z805939-1EA UN3266, Hellmanex III - Hellma Analytics)

NOA 81 (Norland Optical Adhesive 81, Norland Products INC, PN 8101)

##### 1.3.b. Equipment

Compressed air

UV lamp (UV KUB2, KLOE)

Oven or heating plate (from 60°C to 120°C)

Sonicating bath (AL 04-30, Advantage Lab)

### 2. Mounting of the flow chamber and passivation of the microwells

#### 2.1. Passivation of microwells

##### 2.1.a. Material

Slides (Superfrost Microscope Slides LR90SF02, Thermoscientific Menzel Gläser)

SilanePEG 30k Da, PSB-2014, Creative PEG works

HCl 37%

EggPC /L-α-phosphatidylcholine (Egg, Chicken) (840051C-25mg, Avanti)

ATTO 647N labeled DOPE (AD 647N-161 deshydrated, ATTO-TEC). MW: 1485g/mol

BSA (dissolved at 10% in water) (A7030, Sigma, CAS 9048-46-8)

Pluronic F-127 (P2443, Sigma, CAS 9003-11-6)

Chloroform 99% (32211, Honeywell, CAS 67-66-3)

Methanol 99.9% (34860, Sigma, CAS 67-56-1)

Ethanol 96% (UN1170, Carlo Erba, CAS 64-17-5)

Kimwipes

##### 2.1.b. Equipment

Compressed air

Plasma cleaner (air). Plasma DIENER electronic ZEPTO

Nitrogen

Hamilton syringes (volume between 1 and 100 uL)

Sonicator (micro-probe) (Vibracell 72343, Bioblock Scientific)

### 3. Filling of the microwells with reaction mix and closing of the microwells

#### 3.1. Filling of the microwells

##### 3.1.a. Material

Reaction mix with proteins (home-made purified or bought from cytoskeleton inc. for example)

Kimwipes

#### 3.2. Closing of the microwells

##### 3.2.a. Material

Mineral oil (RTM13 Viscosity Reference Standard, Paragon scientific)

VALAP (1/1/1 w/w/w; Vaseline, 5775.1, Carl Roth; Lanoline, 5313.1, Carl Roth; Parafin, 26157.291, VWR)

Kimwipes

## Method

### 1. Microfabrication of SU8 and PDMS molds in order to build the microwells

#### 1.1. SU8 mold fabrication

##### 1.1.1. Method (Figure 1A)

◯ <u>Microwell design</u>. Design should be first drawn with the desired dimensions in microns on Inskape software for example if using a PRIMO system (Alveole), converted into a pdf format with structures in white on a black background. Design can also be drawn on 8-bit image of 1824 × 1140 px^2^ (Fiji).
◯ <u>Spincoating.</u> Spin-coat clean glass wafers with SU-8 (3025 or 3050 depending on the desired microwell height) following manufacturer’s recommendations, to obtain the desired thickness, and soft-bake: first at 65°C and then increase the temperature by 5°C every minute to reach 95°C.
◯ <u>UV Exposure.</u> Position the wafers on the inverted microscope equipped with PRIMO system using the desired magnification. Load the PDF images into the LEONARDO software and project them onto the resists at a dose following recommendations of SU-8 manufacturer and Alveole. In the case the final UV pattern to be projected is bigger than the field of view of the objective, the image is automatically stitched and the final structure is obtained by digitally-controlled sequential aligned UV illumination, allowing for the formation of watertight channel.
◯ <u>UV Exposure (alternative).</u> Alternatively for the exposure step, a quartz photomask or a printed mask (lower resolution) can be used with a UV exposer (KUB2 - MANUFACTURER or MJB4 - MANUFACTURER) following SU-8 manufacturer’s recommendations.
◯ <u>Post Exposure Baking.</u> Place the substrates on a hot plate at 65°C and increase the temperature by 5°C every minute to reach 95°C before a final 8 minutes at 95°C.
◯ <u>Development.</u> Develop the structured SU-8 under agitation for 30 min.
◯ <u>Hard Baking.</u> Bake the substrate at 150°C for 2 hours. *NB. Baking and development times are indicated for a typical height of 25 μm using SU8-3025. For different heights and types of SU8, these times must be adapted according to the manufacturer’s recommendations*.

**Figure 1.**
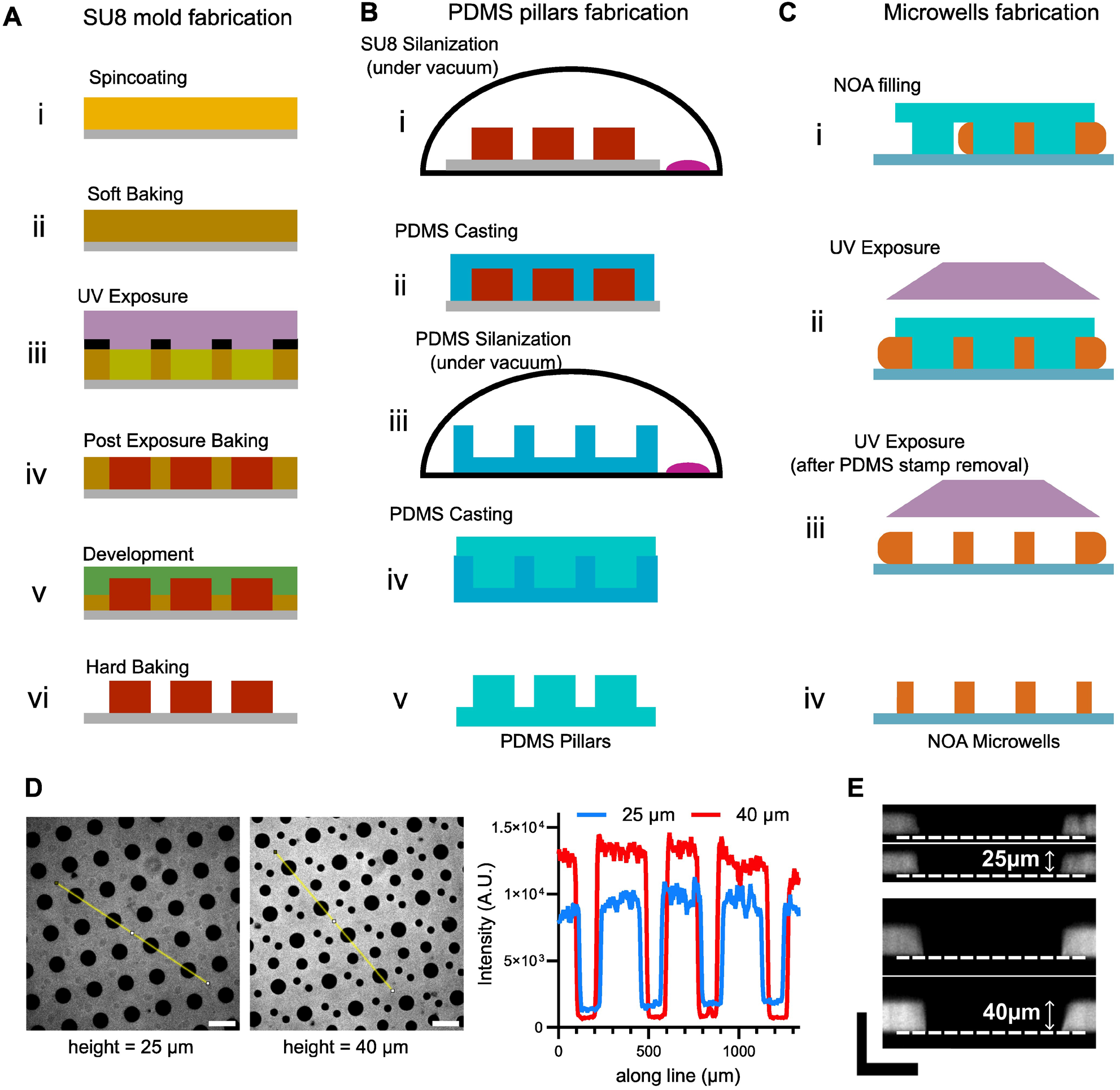
Fabrication of NOA microwells and characterization of their height. **A**. Schemes for the fabrication of SU8 mold. **B**. Schemes for the fabrication of PDMS pillars. **C**. Schemes for the microwells fabrication. **D**. Left: snapshots of NOA microwells with different heights. Right: fluorescence profiles (taken on the yellow lines in the images) of the microwells. Scale bar is 200 μm. **E**. Resliced z-stacks of microwells to measure microwell height. Scale bars are 50 μm.

#### 1.2. Primary PDMS mold and PDMS pillars fabrication

##### 1.2.1. Method (Figure 1B)

◯ <u>SU8 Silanization.</u> Silanize the SU8 mold with the following protocol: place the sample and a drop of 5-10μl of silane onto a glass cap in the glass desiccator. Make the vacuum 5 min then close the valve, switch off the pump and let react for 1 hour. Open the valve then the desiccator and bake samples at 100-150°C for 2 hours.
◯ <u>PDMS Casting.</u> Take the SU8 mold and pour PDMS (previously degassed) on top of it. Polymerize at 70°C overnight.
◯ <u>PDMS Silanization.</u> The next day, take the PDMS primary mold, place it in a glass Petri dish and silanize it according to the protocol described above, with the following modification: bake at 70°C to avoid vitrification of the PDMS.
◯ <u>PDMS Casting.</u> Pour PDMS (previously degassed) on top of the silanized PDMS primary mold to obtain the PDMS stamps necessary for microwells fabrication.
◯ <u>Epoxy mold.</u> At this stage, the PDMS pillars obtained in the previous step (step V, figure 2B) can be used to produce an epoxy resin mold. The advantage of an epoxy mold is that it can be reused more frequently than the primary PDMS mold and does not require silanization. The protocol for preparing the epoxy mold is as follows: prepare the epoxy resin with a weight ratio of 100 g resin:50 g hardener, and mix the two components thoroughly. Centrifuge for 5 minutes at 500 g to eliminate air bubbles. Pour the epoxy into the bottom of a glass Petri dish. During epoxy resin preparation, the PDMS piece can be placed under vacuum. This will reduce air bubbles that may form when the PDMS comes into contact with the epoxy resin. Then place the PDMS piece directly on the epoxy, pillars down (it should be brought at an angle into the epoxy solution, to avoid air bubbles). The PDMS will float on the epoxy. Wait 48 hours at room temperature for the resin to harden. Finally, remove the original PDMS and recast new PDMS onto the epoxy mold to create new pillars.

**Figure 2.**
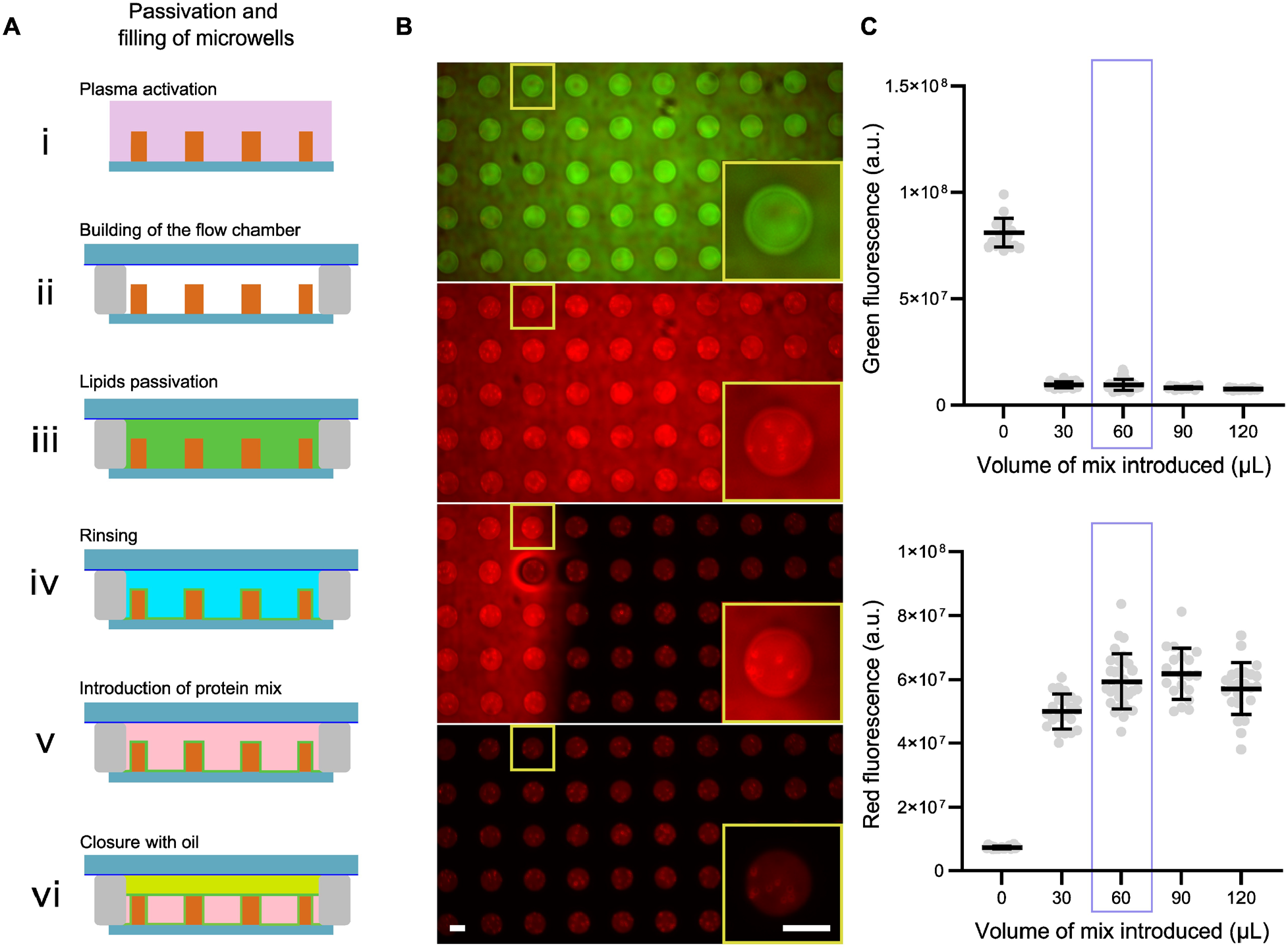
Filling and closing of the microwells. **A**. Schemes for the passivation, filling and closing of microwells. **B**. Images showing microwells first filled with a green dye, then replaced with a red dye and closed with mineral oil. Scale bars are 50 μm. **C**. Estimation of the optimal volume of reaction mix to introduce in microwells.

#### 1.3. NOA microwells preparation

##### 1.3.1. Method (Figure 1C)

###### 1.3.1.1. Cleaning of the coverslips

◯ Wipe the coverslips with Ethanol 96%
◯ Rinse with mqH_2_O
◯ Sonicate for 30 minutes at 60°C in Hellmanex 2%
◯ Rinse by putting the holder with the coverslips in a big volume of mq water (2 L) with agitation. Leave the coverslips for at least 30 minutes in the water.
◯ Dry with compressed air before using

###### 1.3.1.2. Filling the NOA under the PDMS stamps

◯ <u>Stamp preparation.</u> From the PDMS mold prepared in the previous section (1.2), remove the whole PDMS stamps. Cut the rectangle containing the PDMS stamps. Then cut the PDMS in small pieces (about 0.6 cm x 1 cm) that will be the stamps used for microwell fabrication. The cutting should be done in one time (single scalpel cut, sharp cut). Clean the stamp with compressed air and put it on a clean coverslip. Press a little bit on it until you don’t see air in-between the glass and the PDMS. *NB. The PDMS primary mold should not be used more than 4 or 5 times. After that, the microwells are deformed. For a longer-lasting mold, prepare an epoxy mold (see section 1.2.1)*.
◯ <u>NOA Filling.</u> Put a drop of NOA on one side of the stamp and wait for the NOA to diffuse by capillarity around the pillars. When the NOA went under the whole stamp, remove the excess of NOA by aspirating with a pipette
◯ <u>UV Exposure.</u> Expose the NOA to UV lamp for 12 minutes at maximum power. After exposure, cut with the scalpel to remove the NOA around the wells. The cuts should be straight.
◯ <u>UV Exposure (2).</u> After the cutting, expose again the NOA to the UV for 2 minutes at maximum power
◯ <u>NOA Curing.</u> Cure the NOA on a hot plate (110-115°C) for at least 2 hours or at 60°C overnight. After this step, microwells can be stored at room temperature for many days/weeks.

##### 1.3.2. Characterization

As NOA can be cross-linked using UV light, it remains auto fluorescent at these wavelengths. This effect can be attenuated after long exposure to UV light. We take advantage of this intrinsic property to check microwell quality. We use this property to check microwell quality. Firstly, by visual inspection of an image in the DAPI channel, the microwell background should be black, compared with the surrounding microwell walls. In addition, examination of the fluorescence profiles (**Figure 1D**) enables the relative height of the microwells to be estimated. For a more accurate estimate, a z-stack of the microwells can be taken with a confocal microscope. Microwell height can then be measured from a reslice of the z-stack (**Figure 1E**).

### 2. Mounting of the flow chamber and passivation of the microwells

#### 2.1. Passivation of microwells with lipids

This is our preferred configuration at the moment.

##### 2.1.1. Method (Figure 2A)

◯ Rinse (with ethanol then mqH_2_O) then dry a Silane-PEG coated slide, and keep it in a clean box.
◯ Prepare the double tape (180 μm in height) that will come on the two sides of the stamp. Cut the tape precisely so that there is no gap between the stamp and the double side tape (but the tape should not be on the wells).
◯ <u>Plasma activation.</u> Plasma clean the coverslip with microwells for 2 minutes at 80% plasma power.
◯ <u>Building of the flow chamber.</u> Directly after the plasma (be quick), put the scotch tapes on the coverslip and bond the coverslip to the SilanePEG-coated slide.
◯ <u>Lipid passivation.</u> Insert directly the lipids mix (see protocol in (Colin *et al*, 2023b) for lipid mix preparation) in the chamber (about 30 μL). Insert with the pipette on one side and absorb with a kimwipe on the other side (to make the flow). Turn upside down the chamber and hit gently the bottom of the chamber to make sure that the bubbles are leaving the microwells (you can redo one passage of lipids mix to make sure that the bubbles are not in the chamber anymore). Incubate the flow chamber for 10 minutes at room temperature (make sure that the flow cell doesn’t dry during the passivation).
◯ <u>Rinsing.</u> Rinse with 5*200 μL of SUV buffer (10 mM Tris pH 7.4, 150 mM NaCl, 2mM CaCl_2_) and then with 200 μL of the buffer of the experiment.

##### 2.2. Passivation of microwells with BSA/Pluronic

◯ Follow the same steps as describe in the previous section but instead of passivating with lipids, passivate with a solution of BSA and Pluronic F-127 (1% final concentration, in water, filtered).
◯ Rinse with 5*200 μL of mqH_2_O and then with 200 μL of the buffer of the experiment.

##### 2.3. Passivation of microwells with SilanePEG

◯ Plasma clean several coverslips with microwells for 5 minutes at 80% plasma power.
◯ Incubate them in a SilanePEG 30k solution with agitation overnight. (N.B: do not warm the silane solution before use).
◯ After the overnight coating, wash the glasses with ethanol then with water.
◯ Dry with compressed air.

### 3. Filling of the microwells with reaction mix and closing of the microwells

#### 3.1. Methods

◯ <u>Introduction of protein mix.</u> Introduce 60 μL of protein mixture into the chamber. If the mixture contains solid objects (e.g. beads), turn the chamber upside down for 3-5 minutes to allow the objects to settle.

#### 3.2. Characterization (Figure 2B, 2C)

To ensure that the desired amount of protein is introduced into the microwells, the volume of the reaction mixture must be carefully determined. To visualize the volume required to fill the microwells, we performed the following experiment: we first filled the microwells with a green dye. We then replaced this green dye with a mixture of red dye (in volumes ranging from 0 to 120 μL). After closing the microwells, we measured green and red fluorescence for the different volumes of red dye introduced (**Figure 2C**). We observed that a volume of 60 μL is required to completely replace the green dye with red dye. We therefore decided to use a volume of 60 μL for the reaction mixture.

*NB: this optimal volume will vary if the height of the flow chamber is modified*.

#### 3.3. Closing of the microwells

##### 3.3.1. Methods

###### Closure with oil

Close microwells by introducing 50 μl of mineral oil into the flow chamber. Use a minimum flow rate to close the microwells slowly. If the oil is introduced too quickly, it will tend to enter the microwells more deeply, creating a larger meniscus.

##### 3.3.2. Characterization

###### Control of microwells closure

The microwells can be used for experiments on very long time scales (several hours). However, in order to be sure that the amount of proteins stays the same (and limited) as a function of the time, we verified that the microwells were well closed, and that they stayed closed for a very long time. For that, we bleached the full fluorescence present in the microwell and we measured the recovery of fluorescence as a function of time (**Figure 3A**). We observed and quantified that the microwells stay closed for several hours. In comparison, when the same kind of FRAP is done on an open well, the fluorescence is recovered in few minutes, showing a constant exchange of the microwells with the outside medium (**Figure 3A**).

**Figure 3.**
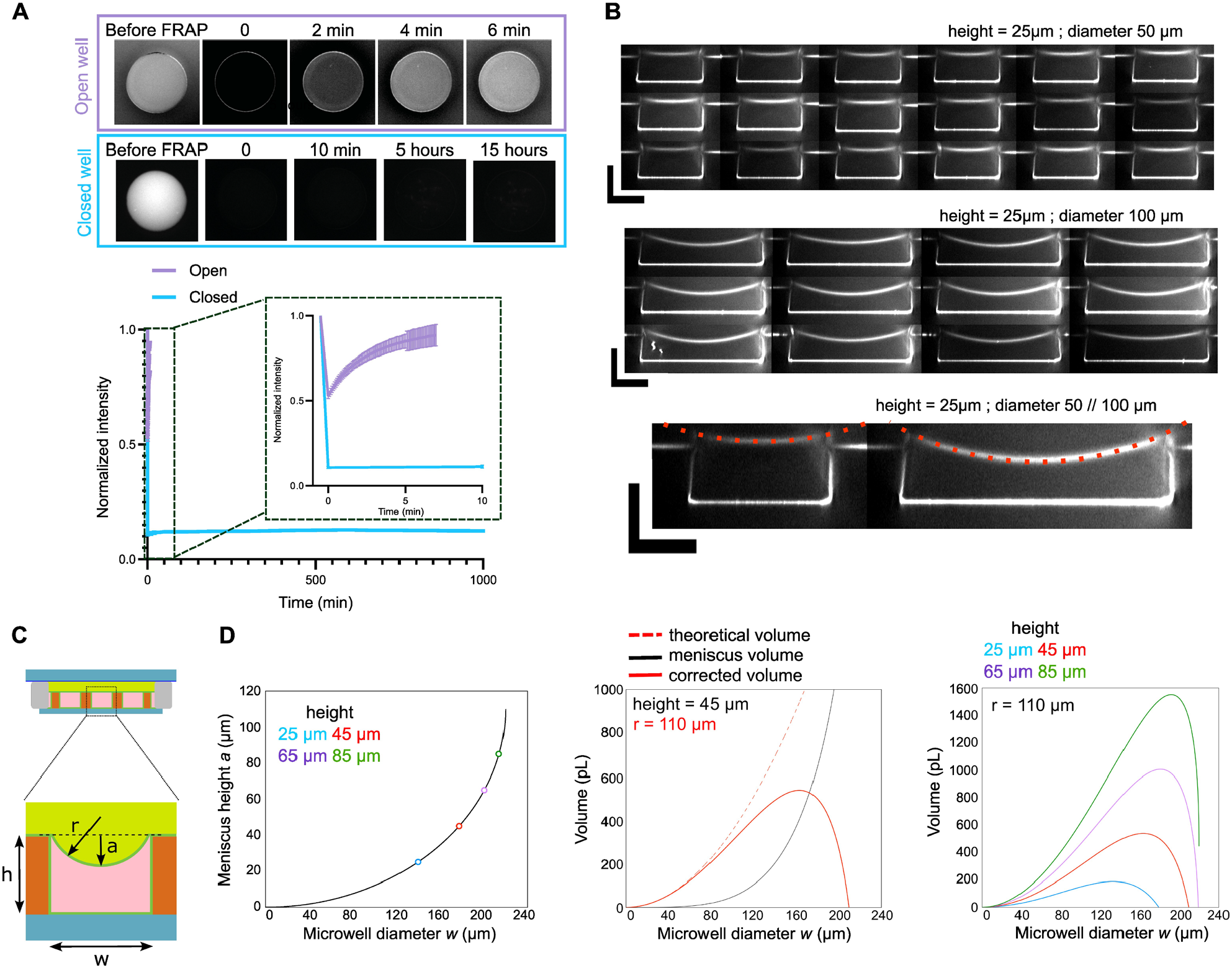
Control of microwell closure. **A**. Top: images of open and closed microwells where the fluorescence was bleached and monitored over time (FRAP experiment). Bottom: Curves showing fluorescence recovery as a function of time for open and closed microwells. **B**. Examples of resliced z-stacks for microwells with diameters of 50 or 100 μm and heights of 25 μm. Curvature radius (r = 110 μm) was measured on those images and was similar for both diameters. Scale bars are 25 μm. **C**. Scheme of the microwell with the meniscus. **D**. Left: Estimation of meniscus height as a function of microwell diameter. Colored circles represent maximum possible microwell diameter for various microwell heights. Middle: Theoretical and corrected microwell volume as a function of microwell diameter. Volume was corrected with meniscus height. Right: Corrected volume as a function of microwell diameter for various microwell heights.

###### Meniscus and geometry: correction of the theoretical volume

By doing z-stacks on microwells with different diameters, we observed that microwells have a more prominent meniscus (**Figure 3B**). Since precise volume of the microwell is important to estimate the number of molecules present in the microwell, we estimated the volume loss caused by the meniscus with the formula below (**Figure 3C, 3D**). These curves can then be used to correct the theoretical volume of the microwell.

In addition, as it can be seen on the curve in **Figure 3D**, for a given height, if the diameter of the microwell is too big, the meniscus volume will be higher than the microwell volume. Therefore, from this curve, it can be seen that there is a maximum diameter of microwell for a given height. Above this maximum diameter, the microwell will be filled by oil and not anymore by the protein mix.

###### Equations for the correction of theoretical volume with meniscus volume

*a* meniscus height / *h* microwell height / *w* microwell diameter / *r* curvature radius

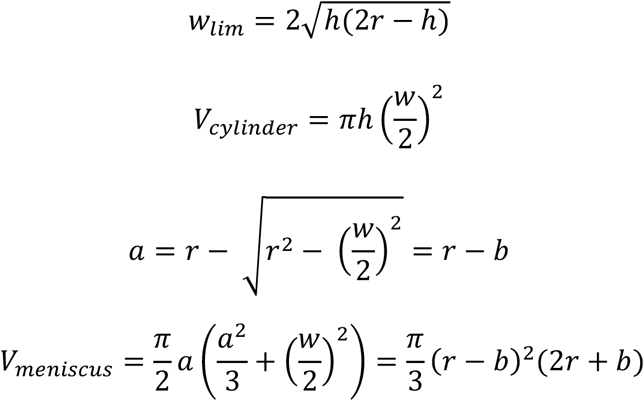

## Results

### Generation of microwells with various sizes and shapes

With the initial design, several sizes and shapes can be drawn (section 1.1). The only limitation is that the well design must be full (i.e. no additional structures can be located inside the microwell). This is because the NOA flows around the shape of the pillars, therefore any hole inside the pillar cannot be filled by the NOA. This makes it possible to generate microwells with perfectly controlled volumes and surface-to-volume ratios, as long as the maximum microwell characteristic side is not exceeded (see section 3.3).

The different sizes and shapes obtained (**Figure 4A**) can then be used to vary the amount of components in the microwell in a controlled way. By comparing structures formed in wells of the same volume but with a different shape, the decoupling of mechanical stress from the quantity of components can also be studied.

**Figure 4.**
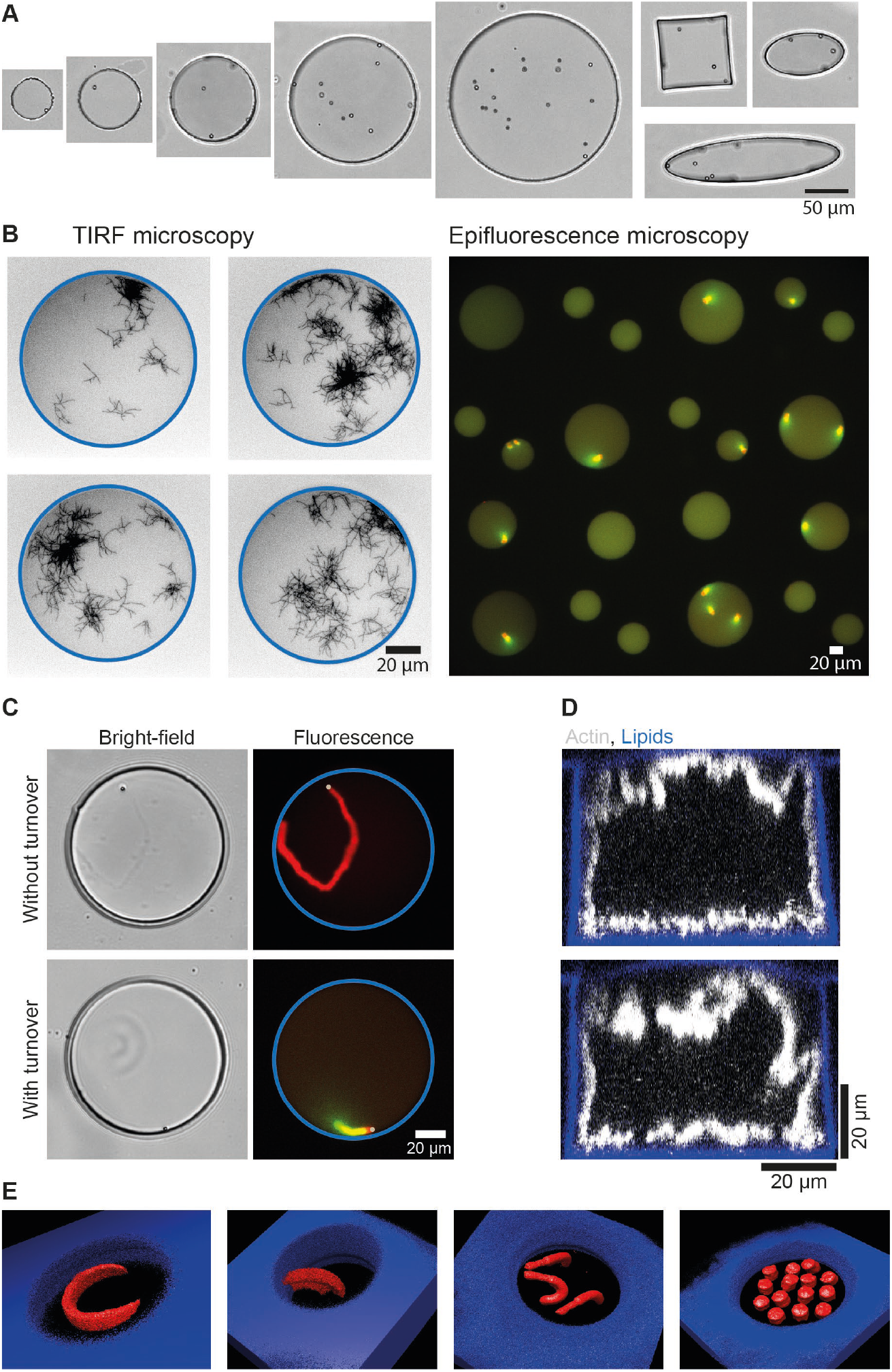
Examples of imaging and functionalization of microwells in order to study actin-related processes. **A**. Bright-field images of microwells with different sizes (from 50 to 200 μm in diameter) and with different shapes (circle, square or ellipse). **B**. Imaging. Left: images of microwells with branched actin network observed with TIRF microscopy. Right: array of microwells imaged with epifluorescence microscopy (20X objective). **C**. Introduction of polystyrene beads (2 μm diameter) in microwells for the reconstitution of actin structures without or with turnover. **D**. Reconstruction of confocal z-stacks showing the growth of actin network from microwells walls from biotin-lipids functionalized with a streptavidin nucleation promoting factor for branched actin networks. **E**. Micropatterning at bottom of microwells. 3D reconstructions of various micropatterns designs leading to various actin network shapes.

### Microwells are adapted to several types of microscopies

For imaging, one of the main advantages of microwells is that they have a glass bottom, making them suitable for several types of fluorescence microscopy. Indeed, single actin filaments can be visualized with Total Internal Reflection Fluorescence microscopy (TIRF) (**Figure 4B**), z-stacks of the entire microwell volume can be taken (**Figure 1E, 4D, 4E** (Yamamoto *et al*, 2022; Sciortino *et al*, 2025)), and large fields of view can be taken with epifluorescence microscopy, making it possible to generate large amounts of data from microwell experiments (Guérin *et al*, 2025). This makes microwells a versatile platform, suitable for many types of fluorescence imaging.

### Introduction of micron-sized beads in microwells

As mentioned in section 3.1, solid objects can be introduced easily in microwells. For example, micron-sized beads to study actin-based motility could be introduced in the microwells (**Figure 4C**). Those beads are coated with a protein promoting the growth of branched actin networks. Therefore, introduction of beads in microwells allowed for the determination of minimal conditions needed to maintain actin networks in a dynamic steady state (based on assembly, disassembly, and recycling) that generated bead movement (Colin et al, 2023a). In a similar setup, protein turnover was demonstrated to be essential for the coexistence of several actin networks that compete for a limited pool of proteins (Guérin *et al*, 2025).

### Microwells walls can be functionalized

As explained in section 2.1, the microwells are currently passivated with lipids. For simple passivation, non-functionalized lipids can be used. For some experiments, growth of an actin network from the microwell walls may be desired. In such cases, functionalized lipids (with biotin or nickel) can be used. Since the lipids cover all sides of the microwells, including the interface with oil, actin network growth can be observed from all sides when functionalized lipids are employed (Yamamoto *et al*, 2022; Sciortino *et al*, 2025), **Figure 4D**.

### Micropatterning at the microwell bottom

When microwells are passivated with SilanePEG (section 2.3), it is then possible to create lipid micropatterns at their bottom (**Figure 4E**, (Guérin *et al*, 2025)). To achieve this, the micropattern design is entered into the Leonardo software that controls the Primo device. As shown in **Figure 4E**, several shapes can be drawn, enabling actin networks with varied architectures to be obtained, developing in three dimensions as a function of the initial biochemical conditions. In addition, by adjusting the gray level of the input pattern, it is possible to have different protein densities on the pattern and thus generate actin networks with different densities (Guérin *et al*, 2025).

## Discussion

In this protocol, we explain how to prepare NOA microwells. With the exception of the initial manufacture of the SU8 mold, the microfabrication of NOA microwells does not require sophisticated or expensive equipment. The microwells can be prepared in advance and stored for a few weeks, limiting preparation time on the day of the experiment. They are therefore easy to handle and use on a daily basis. Sizes and shapes can be extensively varied depending on the research questions, provided the ratio between microwell diameter and height remains within appropriate limits (see section 3.3.2).

The versatility of microwells enables their compatibility with numerous fluorescence microscopy techniques. Furthermore, they facilitate the generation of large datasets (up to 300 wells per experiment). Additionally, imaging microwells with an objective having sufficient working distance enables quantitative estimation of fluorescent protein amounts in both cytoskeletal structures and the microwell bulk. This allows accurate determination of depletion effects occurring within these small volumes (Colin *et al*, 2023a; Guérin *et al*, 2025).

If compared with other methods of creating confinement for reconstituted cytoskeleton networks, one significant limitation is that microwell shapes cannot change during experiments, unlike water droplets in oil or GUVs. However, because microwells are easily manipulated, they can first be used to determine the optimal biochemical conditions for observing desired phenomena, with findings subsequently transferred to deformable compartments.

Another limitation of microwells is that their protein content cannot be modified during experimentation. The possibility of perturbing microwells during the course of an experiment would make it possible to study structural adaptability in response to perturbations, or to counteract the effects of protein aging, which appear after several hours (Colin et al, 2023a). While the current configuration does not allow for such perturbations, future methodological developments will likely incorporate this important capability.

## Acknowledgments

This work was supported by the European Research Council (consolidator grant 771599 [ICEBERG] to M.T. and advanced grant 741773 [AAA] to L.B.) and the grant ANR-24-CE13-3582 SCALING to A.C. This work was also supported by the MuLife imaging facility, which is funded by GRAL, a program from the Chemistry Biology Health Graduate School of University Grenoble Alpes (ANR-17-EURE-0003).

